# Multi-Species, Genome-Wide Metabolic Network Reconstructions Reveal the Basis for Metabolic Versatility in Mycobacteria

**DOI:** 10.64898/2026.09.02.748904

**Authors:** Ignacia Cancino Aguirre, Manisha Priya, Acely Garza-Garcia, Luiz Pedro S. de Carvalho, Hidde de Jong, Delphine Ropers

## Abstract

The genus *Mycobacterium* comprises over 200 species, many of which now have complete genome sequences. Some are major pathogens causing diseases like tuberculosis and leprosy, while others are harmless environmental organisms with useful abilities such as degrading pollutants. Environmental mycobacteria are often seen as metabolic generalists, able to utilise a wider range of carbon sources than host-associated species, which are typically more specialised due to their restricted habitats. This metabolic versatility has been proposed to stem from differences in nutrient uptake capabilities rather than catabolic pathways. In order to test this explanation, we developed and validated genome-scale metabolic models for five *Mycobacterium* species with varying lifestyles and growth rates, creating a computational approach enabled by CarveMe that allows rapid construction of models from genome information. By combining these models with microbiology experiments the study showed that the capacity of the bacteria to transport nutrients into the cell is indeed key to metabolic versatility. We notably found through load-partition experiments that, if a transporter is present but cannot take up its substrate at a rate sufficient for growth, the supply of multiple substrates can mitigate this rate-limiting step. This suggests that mycobacterial species have evolved high-affinity, low-rate systems for nutrient uptake in their ecological niches. More generally, our results demonstrate that a combination of automated annotation methods and straightforward bacterial physiology experiments allow the reconstruction of metabolic models of good predictive quality for hitherto little studied mycobacterial species.

## Introduction

The genus *Mycobacterium* comprises more than two hundred recognized species, many of which now have complete genome sequences (***Tortoli, 2014; Bachmann et al., 2020; Armstrong et al., 2023***). Some mycobacterial species are important human and animal pathogens, including *M. tuberculosis, M. bovis* and *M. leprae*, causal agents of human and bovine tuberculosis and leprosy, respectively. Others are environmental, non-pathogenic organisms, with important biochemical abilities, including degradation of hydrocarbons, thermoplastic polymers such as polyethylene terephthalates or polycyclic aromatic hydrocarbons. Furthermore, an increasing number of infections with opportunistic pathogenic species have been observed. For example, *M. abscessus* is an important pathogen in the context of cystic fibrosis and patients with certain types of lung disease (***Degiacomi et al., 2019; Brugha and Spencer, 2021***). Although still in small numbers, compared to diseases such as tuberculosis and leprosy, infections with these non-tubercular mycobacteria (NTM) are increasingly prevalent and have their own unique challenges associated with antibiotic treatments, absence of vaccines or ideal diagnostics (***Bryant et al., 2016; Baldwin et al., 2019***).

Environmental mycobacteria have been considered capable of growing on a broader range of carbon sources than host-associated species (***Edson, 1951***). This notion stems from the fact that environmental species are subjected to a wider variety of conditions and nutrients in the environment, and therefore must be generalists, in contrast with obligate pathogens, that are only found inside a host or a set of hosts, and hence, are more specialized. An initial bioinformatics analysis comparing the genomes of the human-restricted slow grower *M. tuberculosis* and the environmental fast-grower *M. smegmatis* suggested that this difference in metabolic versatility may be due to the capability to take up nutrients (***Titgemeyer et al., 2007***). In particular, *M. smegmatis* has 28 putative carbohydrate uptake systems, while only five were found in the genome of *M. tuberculosis*. This hypothesis implies that many differences in metabolic capacities may derive from the presence or absence of transporters systems rather than specific catabolic pathways.

This work was designed to test the hypothesis that differences in nutrient utilization in the *Mycobacterium* genus arise from differences in nutrient uptake, and not due to presence or absence of catabolic pathways. To do this, we developed an ensemble of validated genome-scale metabolic models for five distinct *Mycobacterium* species, spanning the entire genus, with different lifestyles and growth rates. By means of our computational approach, enabled by CarveMe (***Machado et al., 2018***), models of any mycobacterial species can be quickly and straightforwardly assembled. We used the genome-wide metabolic models alongside with microbiological experiments to demonstrate that much of mycobacterial growth physiology can indeed be described by differences in nutrient uptake.

## Results

### Distinct growth performance of selected *Mycobacterium* species

In order to investigate the relation between nutrient transport and metabolic versatility of NTM, we selected four, phylogenetically and ecologically diverse *Mycobacterium* species, as well as *M. tuberculosis* for reference (**Fig. 1**). *M. abscessus* is an environmental species that has recently adapted to survive and proliferate within host tissues (***Bryant et al., 2013, 2021***). It thrives in diverse environments, including aquatic environments and host lungs, and has become an emerging threat to global health (***Johansen et al., 2020***). *M. aromaticivorans* is an environmental species that was initially isolated from oil-contaminated soil and water (***Hennessee et al., 2009***). *M. smegmatis* is a non-pathogenic species, which has been widely used as a model for *M. tuberculosis* due to their similar gene content (~75% of protein-coding genes), cell envelope composition, and antibiotic susceptibility (***Sparks et al., 2023***). *M. marinum* is a fish, amphibian, and human pathogen, causing cutaneous infections in mammals. It can also persist independently in specific aquatic habitats (***Canetti et al., 2022; Dong et al., 2021***). **Supplementary Table 1** summarizes essential characteristics of the species considered here.

**Figure 1.**
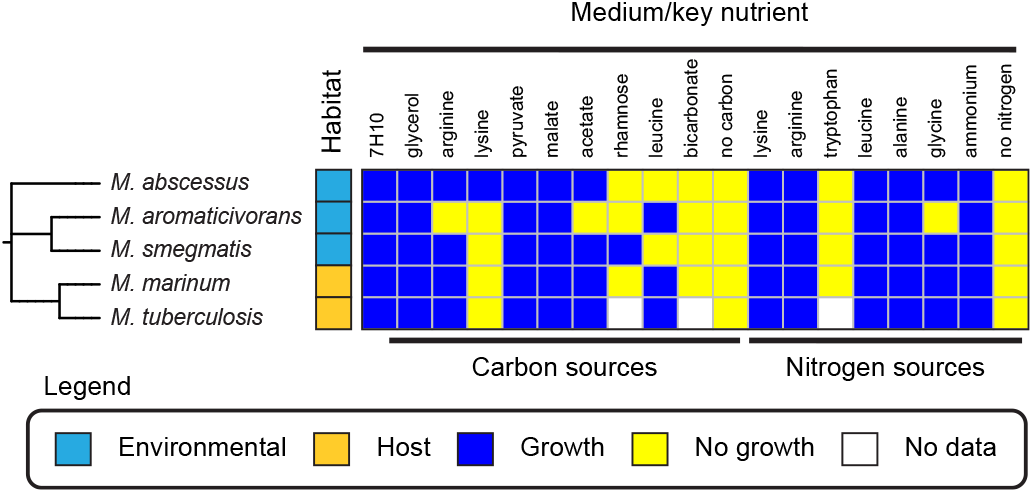
Selected mycobacterial species and their growth on different carbon and nitrogen sources. Phylogenetic relations between the five selected species, their typical habitats, and results of agar plate experiments using Roisin’s minimal medium with a defined carbon or nitrogen source. The data for *M. smegmatis, M. abscessus, M. aromaticivorans* and *M. marinum* are representative from two independent experiments. The data for *M. tuberculosis* are from (***Lofthouse et al., 2013***) (see **Supplementary Fig. 1** for the complete *M. tuberculosis* dataset).

While the nutrient conditions supporting growth of *M. tuberculosis* have been extensively characterized (***Youmans and Youmans, 1953; de Carvalho et al., 2010; Lofthouse et al., 2013***), few systematic studies for the other species exist. Information on growth phenotypes is critical, however, for the automated reconstruction of metabolic networks from annotated genome sequences. We therefore performed growth experiments on agar plates for *M. abscessus, M. aromaticivorans, M. smegmatis* and *M. marinum*, using Roisin’s minimal medium supplemented with a single carbon or nitrogen source (*Methods*). The results are shown in **Fig. 1**. Clear differences between the species can be seen. For example, *M. smegmatis* is the only species capable of growing on rhamnose, while *M. abscessus* is the only species capable of utilizing lysine as sole carbon source. Moreover, every species except *M. aromaticivorans* can grow on arginine and acetate. Literature data on the growth of *M. tuberculosis*, obtained by means of agar plates using the same minimal medium as in our experiments (***Lofthouse et al., 2013***), are shown in **Fig. 1** for reference (see also **Supplementary Fig. 1**). In summary, our data show that NTM display distinct growth profiles, with the four species selected differing in at least one and at most four conditions (out of 16).

### Automated reconstruction of genome-scale metabolic models for mycobacteria

All five *Mycobacterium* species used in this study have been sequenced, but with the exception of *M. tuberculosis* and *M. smegmatis*, little is known about their metabolism. In recent years, powerful methods for the automatic reconstruction of metabolic networks from genome sequences have been developed (***Mendoza et al., 2019***). In this study, we employed a widely-used tool for microbial metabolic network reconstruction, CarveMe (***Machado et al., 2018***). CarveMe is based on a universal genome-scale metabolic model (GSMM) which contains reactions from a variety of organisms. In a top-down approach, CarveMe “carves” a genome-specific model from its universal model based on protein sequence alignment scores and, optionally, a reference GSMM.

As a first step of the application of CarveMe to our network reconstruction problem (**Fig. 2A** and *Methods*), we selected reference genome sequences for the five species considered here. In order to ensure a uniform quality of annotation, we chose EggNOG-mapper (***Cantalapiedra et al., 2021***) to structurally reannotate the genome sequences. Second, we adapted the universal bacterial model of CarveMe to the reconstruction of mycobacterial metabolism. In particular, we improved an existing model for *M. tuberculosis*, iEK1011 2.0 (***López-Agudelo et al., 2020***), among other things by ensuring that a maximum of reactions are chemically balanced and introducing a periplasmic compartment (*Methods*). The reactions of the resulting reference model were added to the universal bacterial model. Third, we modified the code of CarveMe to include new parameters for controlling the reconstruction process. The parameters allow us to more strongly penalize the inclusion of reactions without convincing genomic evidence and reward inclusion of reactions from the *M. tuberculosis* reference model. Fourth, we checked and corrected the network connectivity and mass balances of the draft models generated by CarveMe, using the MEMOTE tool (***Lieven et al., 2020***).

**Figure 2.**
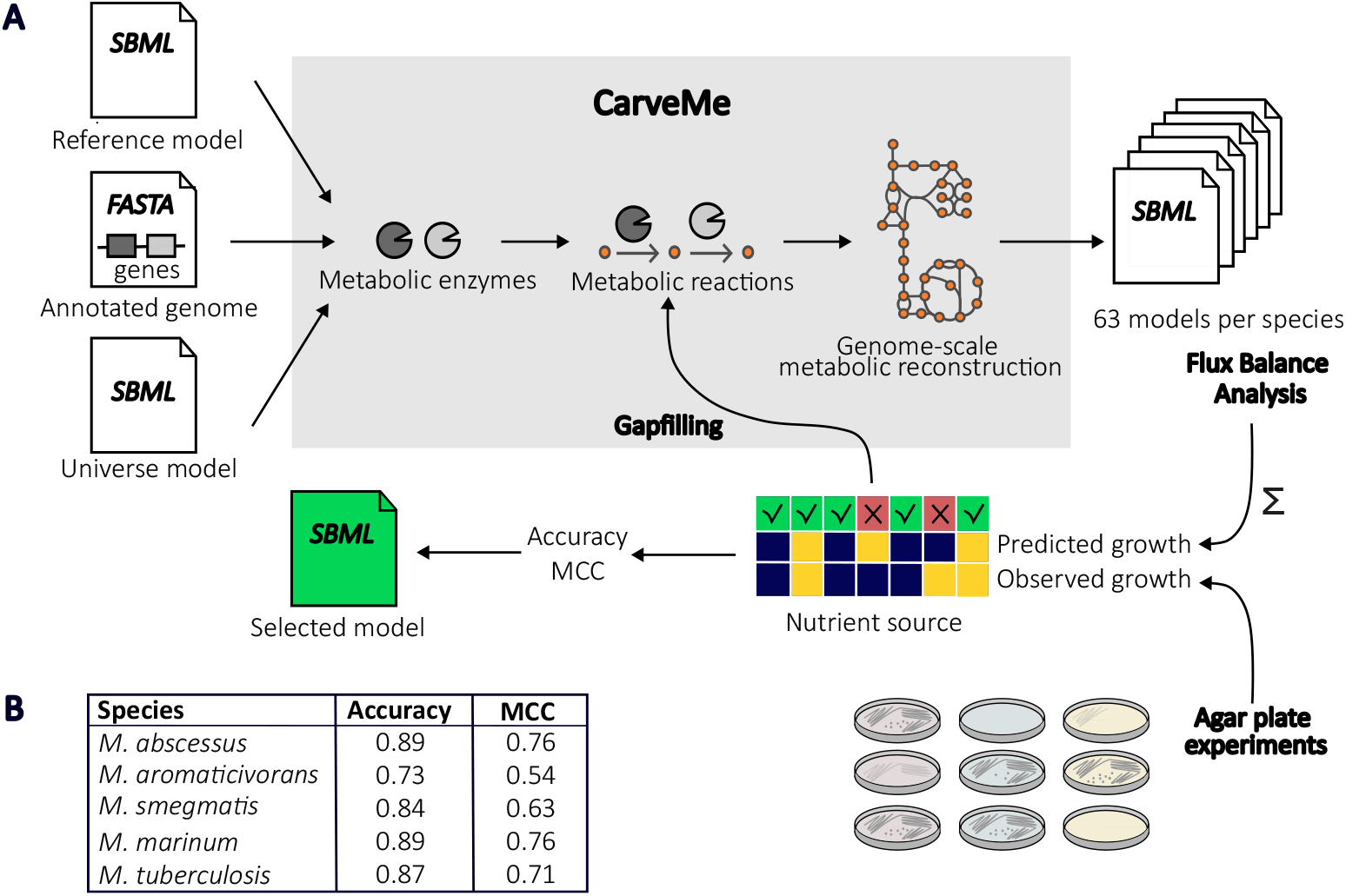
Automated reconstruction of genome-scale metabolic models of *Mycobacterium* species. (A) Outline of the approach, consisting in the generation of GSMMs from genome annotations linking proteins to metabolic reactions in an updated universal model, using a manually curated model for *M. tuberculosis* as reference. The candidate models are tested for consistency with growth data as well as measured nutrient uptake, product secretion, and growth rates. (B) The accuracy and Matthew’s correlation coefficient (MCC) scores for the selected draft models indicate that they have good overall predictive quality.

For different values of the reconstruction parameters, CarveMe generated more than 60 draft GSSMs for each species. In order to check the consistency of the models with the experimental data from **Fig. 1**, we extended the models with the generic biomass and maintenance reactions from CarveMe (***Xavier et al., 2017***). We then used flux balance analysis (FBA) (***Thiele and Palsson, 2010***) to predict whether the network supports growth in the given conditions, where growth corresponds to a non-zero flux through the biomass reaction (*Methods*). We noticed that in the case of *M. smegmatis*, none of the models predicted growth in a few conditions where growth was observed on the agar plates. We corrected for these discrepancies by completing the models with additional reactions allowing growth on the specific carbon or nitrogen source, using the gapfilling option of CarveMe (***Machado et al., 2018***). From the draft models, we selected those that (i) are maximally consistent with the growth data and (ii) have a maximum number of gene-associated metabolic reactions, thus giving high priority to genomic evidence for the reactions.

The correspondence of the selected GSMMs with the growth data, shown in **Supplementary Figs 1 and 2**, is quantified by the accuracy and Matthew’s correlation coefficient (MCC) scores (**Fig. 2B**). While the former refers to the fraction of predictions corresponding to true positives and negatives, the latter expresses the balance between true and false positives and negatives (*Methods*). The accuracy of the models varies from 0.74 for *M. aromaticivorans* to 0.95 for *M. abscessus* and *M. marinum*. The MCC lies between 0.88 and 0.54. The relatively worse performance of *M. aromaticivorans* could be explained by the lower quality of genomic functional annotation for this species (no complete genome available). We also reconstructed the *M. tuberculosis* model and checked that it makes the same predictions as the manually curated reference model.

An example of an interesting result in **Fig. 1** is that, apart from *M. smegmatis*, none of the species grow on rhamnose. The reconstructed models reproduce this observation (**Supplementary Figs 1 and 2**) and allow it to be analyzed in terms of the metabolic capacities of the networks. The network of *M. smegmatis* includes the five key reactions from the consensus pathway for rhamnose catabolism, previously characterized in enterobacteria (***Rodionova et al., 2013***) (**Supplementary Fig. 3**). Alignment of the protein sequences of the enzymes from *Escherichia coli* to the *M. smegmatis* genome shows good correspondence, whereas no or weak matches are found when aligning the sequences to the genomes of the other species.

We also verified whether the reconstructed models are consistent with quantitative information on metabolic fluxes available from the literature. The metabolism of mycobacteria has been significantly less studied at the flux level as compared to model organisms like *E. coli* and yeast, but measurements of chosen uptake, secretion, and growth rates do exist for *M. tuberculosis* and *M. smegmatis* (***Beste et al., 2011; Borah et al., 2021; Hooper, 2008***), and to a lesser extent *M. marinum* (***Dong et al., 2021***). For the species for which data are available, we checked whether the models allow flux distributions satisfying the measured uptake, secretion, and growth rates. As can be seen in **Supplementary Table 2**, the reconstructed models are consistent with the data. While the available data do not cover the entire exometabolome and are therefore less constraining than one might ideally hope for, the correspondence between the models and the data is reassuring.

### Structural and functional differences between reconstructed metabolic networks

Next, we analyzed and compared the structure of the reconstructed networks, connecting any observed differences to known cellular functions of the pathways.

**Fig. 3A** illustrates the size of the best draft GSMMs retained from the reconstruction process (see **Supplementary Table 3**, for further statistics). A first observation is that there are large differences in size between the networks. For example, the reconstructed network for *M. tuberculosis*, which is almost identical to the reference network, has 1501 reactions, whereas the networks for the other species are much larger, up to 3079 reactions for *M. aromaticivorans*. The differences in network size weakly correlate with genome size (*R*^2^ = 0.49, **Fig. 3B**). As expected, environmental species, which need to be able to adapt to a variety of changes, have larger metabolic networks than host-associated species. A second observation is that most reactions in the *M. tuberculosis* network also occur in the other networks and therefore form the core of mycobacterial metabolism. Accessory metabolism, specific to one or more of the other species, adds up to a total number of 1870 reactions.

**Figure 3.**
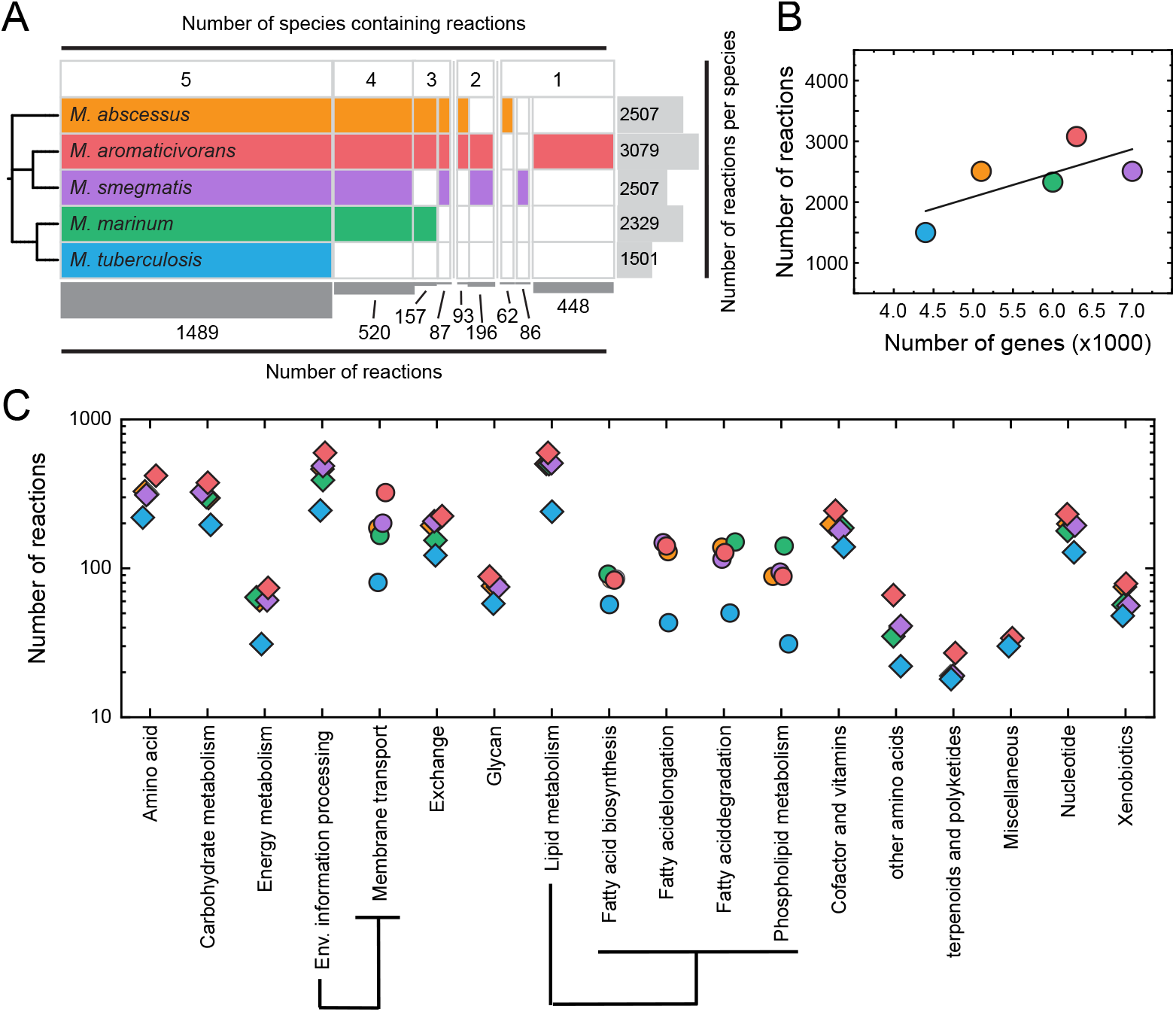
Core and accessory metabolism in the reconstructed models for the *Mycobacterium* species. (A) The summary plot shows the parts of the reconstructed networks shared between species (core metabolism) and specific to each species (accessory metabolism). (B) Identified reactions in the reconstructed networks are weakly correlated with the genome size of the species (*R*^2^ = 0.49). Symbols are data (colors match those from panel A) and the line represents a linear regression of the data. (C) Distribution of the reactions in the reconstructed models over KEGG orthology categories and a few selected subcategories (***Kanehisa et al., 2016***) (colors match those from panel A). Categories with less than 10 reactions are not shown.

The large number of accessory metabolic reactions makes a manual analysis of the differences practically infeasible. We therefore took a coarse-grained, systematic look at the networks by assigning specific metabolic functions to the reactions and testing which functional categories are over-represented in the networks of the individual species as compared to the core network. In order to avoid heterogeneity in existing databases, we reannotated the totality of reactions in our models using the KEGG Orthology (KO) database (***Kanehisa et al., 2016***). We performed Fisher’s exact test for every functional category, with Benjamini-Hochberg correction for multiple testing, to identify statistically significant differences between the networks (*Methods*).

**Fig. 3C** and **Supplementary Fig. 4** show that many of the functional categories distinguishing the metabolic networks of the individual species from the core network of *M. tuberculosis* concern the synthesis and degradation of components of the cell envelope. For example, the reconstructed network of *M. smegmatis* has 284 more reactions in lipid metabolism, confirming conclusions drawn from a previous comparison of the genomes of *M. tuberculosis* and *M. smegmatis* (***Vissa et al., 2009***). The observation agrees with the notion that the cell envelope of *M. smegmatis* has a different and more complex composition and structure than the cell envelope of *M. tuberculosis*, which may limit the use of *M. smegmatis* as a surrogate model for *M. tuberculosis* (***Sparks et al., 2023***). Another functional category over-represented in accessory metabolism concerns the uptake and catabolism of nutrients. This is especially true for the environmental species *M. smegmatis, M. abscessus*, and *M. aromaticivorans*, which is expected as these species have to survive in a more diverse range of environments, containing a higher variety of carbon and nitrogen sources (***Titgemeyer et al., 2007; Niederweis, 2008***).

### Growth differences between species highlight the impact of nutrient transport

How do the structural differences between the metabolic networks of the *Mycobacterium* species translate into growth differences? To answer this question, we chose a dozen different environments, consisting of 7H10 minimal medium supplemented with a specific carbon source of ecological interest for further analysis. Carbon sources include amino acids, nucleosides, monosaccharides, and short-chain fatty acids. We simulated growth of the five species in these environments using FBA and, in parallel, performed agar plate growth experiments. In comparing the model predictions with the data, we took special care in choosing simulation parameters in agreement with the design of the experiment, such as the composition of the medium and the maximum time for colonies to appear (*Methods*).

**Fig. 4A** summarizes the experimental results and the correspondence with the FBA predictions. As expected, the environmental species *M. smegmatis* and *M. aromaticivorans* grow in more conditions (six) than the host-associated species *M. tuberculosis* and *M. marinum* (three). Although *M. abscessus* is an environmental species, it sides with the host-associated species in that it grows in only three conditions. The overall accuracy score of the predictions is 0.65, with errors concentrated on the less well-annotated environmental species. Most errors take the form of false positives, where the models predict growth contrary to what is observed in the experiments. A few false negatives also occur, that is, conditions in which growth is observed but not predicted.

**Figure 4.**
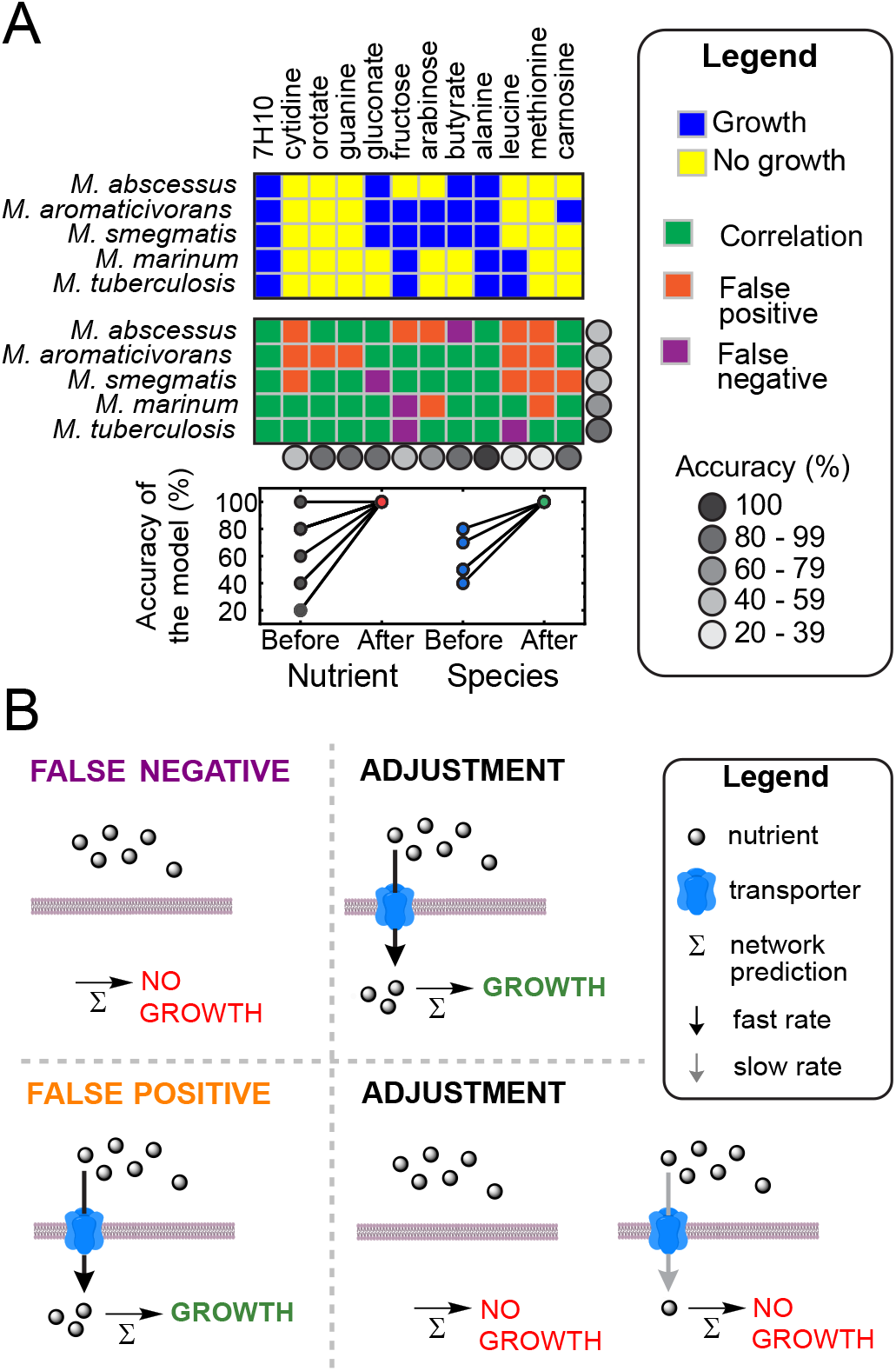
Experimental test of model predictions underlining the role of nutrient transport. (A) Results of agar plate experiments using 7H10 minimal medium with a defined carbon source or 7H10 complete medium as a positive control. (B) Graphical representation of how wrong assumptions on nutrient transport can explain the occurrence of false positives and negatives. The corresponding adjustments make the models completely accurate, as shown in panel A by the statistics for the reconstructed (Before) and adjusted (After) models.

The occurrence of a false negative implies that the metabolic network does not contain reactions for the uptake or utilization of a substrate on which the bacteria are observed to grow. Importantly, we found that a transporter was missing for every false negative encountered in our analysis: growth of *M. smegmatis* on butyrate and gluconate, growth of *M. tuberculosis* on fructose and leucine, and growth of *M. marinum* on fructose (**Supplementary Table 4**). When adding these unknown transporters to the models, and correcting that one of the polyphosphate glucokinase enzymes in *M. tuberculosis* acts not only on glucose-6-phosphate but also on fructose-6-phosphate (***Marrero et al., 2013***), growth was restored for all false negatives. Therefore, insufficient knowledge about nutrient transport systems is the main reason for the occurrence of false negatives in our metabolic reconstructions, not lack of pathways required to break down nutrients (**Fig. 4B**).

False positives occur for two reasons. First, the model erroneously contains a transporter capable of taking up the substrate, or second, the transporter specificity is correct, but the nutrient cannot be taken up at a rate required for growth (**Fig. 4B**). The first case is often the result of annotation errors encountered in the reconstruction process. For example, *M. aromaticivorans* is predicted to grow on guanine, but the model includes a transporter for which there exists only weak evidence: low sequence similarity, just above the 30% threshold, with a suspected transporter from another species (**Supplementary Table 4**). Removing the transporter transforms the prediction into a true negative. Examples of the second case are growth of *M. abscessus* on leucine and fructose. There are no compelling reasons to dismiss the annotation of the transporters for these carbon sources, since sequence similarity with transporters from other species is sufficiently high (55% and 64% identity for leucine and fructose, respectively; **Supplementary Table 4**). When decreasing the maximum uptake rate of the transporters by a factor of 3, however, the predictions are turned into true negatives. While hypothetical, this correction suggests that incomplete knowledge about nutrient uptake kinetics may be responsible for many of the mismatches of the models with the data.

### Kinetic limitations are mitigated by uptake of multiple substrates

Based on the results above, we concluded that the maximum uptake rate for a given carbon source may not be sufficient to sustain growth even when a *bona-fide* transporter exists. There is a straight-forward way to test this hypothesis, which we name “load partitioning”. A fixed amount of carbon can be supplied as a single nutrient or distributed over two nutrients that are independently transported into the cell, splitting the load of carbon uptake. If the maximum capacity of the individual transporters is limiting, one expects that supplying all carbon as a single nutrient cannot support growth, while growth may be possible if the two nutrients are simultaneously taken up through different transporters (**Fig. 5A**). Moreover, it must be possible for the substrates to be co-metabolized together, bypassing diauxic growth. *M. tuberculosis* was the first organism to be shown to be able to co-metabolize carbon sources (***de Carvalho et al., 2010***), therefore enabling this load-partition experiment.

**Figure 5.**
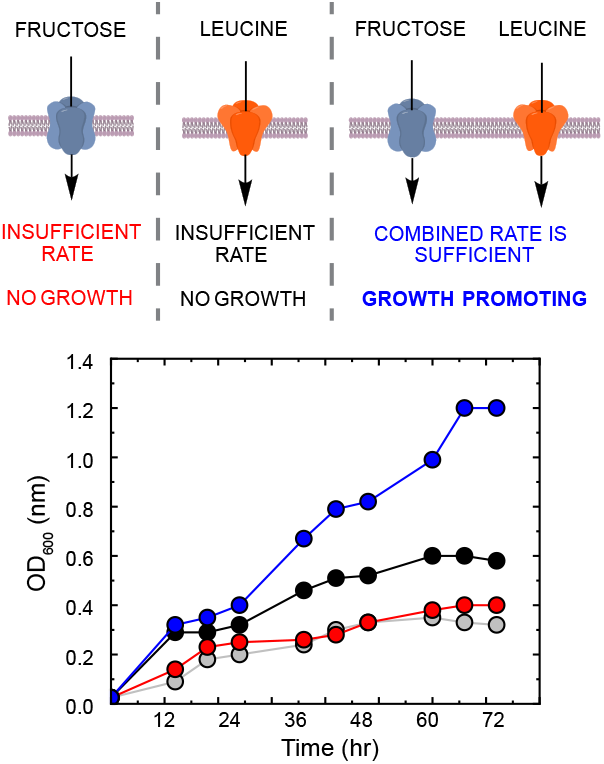
Carbon load-partition experiment for *M. abscessus*. (A) Growth on the individual substrates is not possible due to the limited uptake capacity of their respective transporters. However, the synergy attained through the simultaneous uptake of both substrates provides sufficient resources for cellular growth. (B) Growth curves of *M. abscessus* in 7H9 minimal medium either in the presence of 0.4% fructose (red) or leucine (black), or a combination of the two (blue). As a negative control, the plot also shows residual growth on 7H9 without carbon source (grey).

The data reported above show that *M. abscessus* cannot grow on leucine or fructose as the sole carbon source (**Fig. 4A**). However, the model does contain transporters for these carbon sources, based on sequence similarity with known transporters in other species. This suggests that, when supplying leucine and fructose in combination, *M. abscessus* might be capable of growing. We performed a load-partition experiment, in liquid medium, to test this prediction, by supplying *M. abscessus* with a total concentration of 0.4% of carbon in the form of leucine, fructose, or a combination of the two.

The results of this experiment (**Fig. 4B**) confirm the prediction. Whereas only residual growth, due to the utilization of storage metabolites from the preculture, is observed in the presence of leucine or fructose as the sole carbon source, the culture of *M. abscessus* reaches high biomass levels when the same concentration of carbon is supplied through the combination of leucine and fructose. This observation thus confirms that limitations in transport kinetics can be overcome by the simultaneous uptake of multiple substrates. The experiment also demonstrates that *M. abscessus* is able to co-metabolize carbon sources, similarly to *M. tuberculosis*.

## Discussion

We developed a computational approach for the reconstruction of GSMMs of five representative *Mycobacterium* species, for some of which no models were previously available. We used the CarveMe tool (***Machado et al., 2018***) to reconstruct the models from the genome sequences, supplying it with an improved model of *M. tuberculosis* as a template and the results of agar plate experiments for the growth of the species on selected carbon and nitrogen sources (**Figs. 1** and **2**). A comparison of the reconstructed models revealed that the metabolic networks of environmental species are larger than those of host-associated species, owing in part to the presence of more reactions for the uptake and catabolism of nutrients (**Fig. 3**). Environmental species encounter a broader range of conditions than host-associated species, and are therefore expected to display higher metabolic versatility (***Niederweis, 2008; Cook et al., 2009***).

We validated the reconstructed GSMMs by means of further experiments in which we tested growth on specific carbon sources (**Fig. 4**). The analysis of the results allowed us to confirm a previous suggestion that the difference in metabolic versatility between environmental and host-associated species may be due to the capacity of the bacteria to transport nutrients into the cell, rather than the availability of dedicated catabolic pathways. In particular, in cases where the models do not predict growth when growth was observed (false negatives), the addition of a transporter for the substrate resolved the apparent conflict. This points at a lack of knowledge about transporters for taking up the substrate rather than the availability of metabolic pathways for breaking down the substrate to building blocks for growth. In cases where the models predict growth but no growth was observed (false positives), the discrepancy could often be traced back to low-confidence predictions of transporters. Annotation of membrane transporters is a common issue in metabolic network reconstruction for several reasons, including their lack of experimental characterization, their varied substrate specificity, and their low sequence similarity despite performing the same function (***Casey et al., 2024***).

In a number of cases, there was little doubt about the existence of a transporter, but the bacteria were nevertheless not capable of utilizing the nutrient for growth (**Fig. 4**). This raised the possibility that the maximum uptake rate of the nutrient *via* the transporter may not be sufficient to sustain growth. In order to test this hypothesis, we performed a load-partition experiment for a representative combination of carbon sources (leucine and fructose). We found that, as expected based on our data (**Fig. 3**), *M. abscessus* was not able to grow on either of the carbon sources alone, but did grow when the same amount of carbon was supplied as a combination of the two carbon sources (**Fig. 4**). This scenario matches the conditions generally encountered by mycobacteria in ecological niches, where multiple nutrients are available at low concentrations and co-utilized for growth (***de Carvalho et al., 2010***). Low substrate concentrations put an evolutionary pressure on the development of high-affinity transporters, at the cost of a low uptake rate (***Montaño-Gutierrez et al., 2022***).

The conclusion that nutrient transport is often the rate-limiting step for mycobacterial growth suggests that nutrient transport systems may be potential targets for antibacterial drugs. When inhibiting transporters specific to mycobacterial growth in different infection niches, it might be possible to prevent nutrient uptake and therefore proliferation. Such an approach was previously suggested for *M. tuberculosis* and other pathogenic bacteria (***Davies et al., 2021; Soni et al., 2020***). It notably avoids a problem commonly encountered with antibiotics targeting biosynthetic processes in *Mycobacterium* species, namely the impermeability of the plasma membrane, and efflux systems.

More generally, our results demonstrate that recent advances in bacterial whole-genome sequencing in combination with automated annotation methods and straightforward bacterial physiology experiments allow the reconstruction of metabolic models of good predictive quality for hitherto little studied mycobacterial species. We provide a model reconstruction pipeline, with manually curated annotation resources, that can be readily applied to analyze the metabolism of other mycobacterial species. This is an important asset for the study of novel, poorly characterized NTM species that are a growing concern for human and animal health.

## Methods and Materials

### *Mycobacterium* strains and genome sequences

*M. smegmatis* mc^2^155, *M. abscessus* ATCC 19977, *M. aromaticivorans* JS19b1, *M. tuberculosis* H37Ra, and *M. marinum* ATCC 927 were used in this study. Their genomes, which were used in the model reconstruction process were taken from the NCBI database, with the exception of the genome of *M. marinum*, which was taken from (***Savijoki et al., 2024***). Genome assembly codes and assembly levels are given in **Supplementary Table 5**. All strains used in the growth phenotype experiments were previously genotyped using the housekeeping genes *hsp65, atpA, ffh*, and *recA* as reference.

### Growth experiments

All experiments involving mycobacteria were performed in accordance with BSL-2 biosafety guidelines.

#### Growth of *Mycobacterium* species on selected carbon and nitrogen sources

For model validation (**Fig. 4**), growth on defined carbon sources was assessed using 12-well agar plates. Middlebrook 7H10 medium (Difco BD) was prepared without glycerol. At the time of plate preparation, the 7H10 minimal medium was supplemented with 1× ASC (bovine albumin serum-sodium chloride-catalase supplement; in-house preparation) and 0.2% of a designated carbon source. The following carbon sources were tested: L-methionine, cytidine, gluconate, D-fructose, L-carnosine, L-arabinose, butyrate, L-alanine, L-leucine, guanine, and orotate. One well per plate containing Middlebrook 7H10 supplemented with 0.2% glycerol and 1× OADC (1× ADC, bovine albumin serum-sodium chloride-dextrose-catalase supplement, with oleic acid; BD) served as the positive control. The nutrient sources were from Sigma-Aldrich with the exception of guanine and orotate, which were from Santa Cruz.

*Mycobacterium* species were cultured in Middlebrook 7H9 broth (Difco BD) supplemented with 0.2% glycerol, 1× ADC (BD), and 0.05% tyloxapol (Spectrum). Cultures were incubated at 37°C, or at 30°C for *M. marinum*, with shaking at 150 rpm, until an OD_600_ of approximately 3.0–4.0. Cultures were then washed twice with Middlebrook 7H9 minimal medium (lacking glycerol and ADC), diluted to OD_600_ ≈ 0.05 in medium, and incubated until reaching an OD_600_ of ~0.8–1.0. These secondary cultures were harvested and washed twice with Middlebrook 7H9 minimal medium. The washed cells were diluted 10-fold in Middlebrook 7H9, and 2 *µ*L of the diluted suspension was spotted into each well containing the designated carbon source. Plates were incubated at the temperatures indicated above for 5–7 days for fast-growing species (*M. smegmatis, M. abscessus, M. aromaticivorans*) and 14–21 days for slow-growing species (*M. tuberculosis* H37Ra, *M. marinum*). Growth in each condition was evaluated relative to the positive control.

For the purpose of model reconstruction (**Fig. 2** and **Supplementary Fig. 1**), the above protocol was adapted, so as to allow our data to be combined with datasets from previous work on *M. tuberculosis* (***Lofthouse et al., 2013***). The main change was the use of agar plates with Roisin’s minimal medium instead of Middlebrook 7H10. For conditions of carbon limitation, we added one of the following carbon sources at a concentration of 0.5% (w/v): L-arginine, L-lysine, pyruvate, L-malate, acetate, L-rhamnose, bicarbonate, or L-leucine. NH_4_Cl was added as nitrogen source at a concentration of 0.59%. For conditions of nitrogen limitation, we added one of the following nitrogen sources at a concentration of 0.59%: L-arginine, L-lysine, L-tryptophane, L-glycine, ammonium, L-alanine, and L-leucine. In addition, the medium was complemented with the carbon sources glucose and glycerol at concentrations of 0.5%. As a negative control, one condition had no carbon or no nitrogen source added. As positive controls, we used both Middlebrook 7H10 plates (Difco BD) and Roisin’s minimal media with glycerol, glucose and L-glutamate as carbon sources (3G). The nutrient sources were from Merck.

#### Growth kinetics of *M. abscessus* in load-partitioning experiment

To assess whether poor utilisation of specific carbon sources could be enhanced by supplementation with a secondary carbon or nitrogen substrate, we compared the growth kinetics of *M. abscessus* in the presence of D-fructose, L-leucine, and an equal combination of both (**Fig. 5**). Each condition was tested at a final concentration of 0.4%. The medium was prepared by adding 0.05% tyloxapol (Spectrum) to Middlebrook 7H9 minimal medium (Difco BD). Immediately before use, the medium was further supplemented with 1× ASC and either D-fructose, L-leucine, or a 1:1 mixture of the two at the indicated concentration. Medium supplemented instead with 0.2% glycerol and 1× ADC served as the positive control, whereas medium without supplements served as the negative control.

A preculture of *M. abscessus* was grown to an OD_600_ of ~3.0–4.0, washed twice in the medium lacking carbon sources, and diluted to an OD_600_ of ~0.025 in falcon tubes containing the respective conditions. Cultures were incubated at 37°C with shaking at 150 rpm, and OD_600_ measurements were recorded every 6–12 h to monitor growth kinetics. Growth curves were plotted using Origin-Pro software.

### Reconstruction of genome-scale metabolic models

The reconstruction of the genome-scale metabolic models was performed using the Python-based tool CarveMe (carveme v1.5.1) with IBM ILOG CPLEX studio v22.1 solver (***Machado et al., 2018***). We provided CarveMe with the following inputs: (i) genome sequences for the five species considered here (**Supplementary Table 5**), structurally annotated by means of Prodigal through EggNOG-mapper v2 (***Hyatt et al., 2010; Cantalapiedra et al., 2021***), (ii) a universal bacterial model built from metabolic reactions and metabolites in the BiGG database (***King et al., 2016***), and (iii) a reference model of the metabolic network of *M. tuberculosis* H37Rv used as a template for model reconstruction. The reference model derives from the iEK1011 2.0 model (***López-Agudelo et al., 2020***), which was improved by ensuring maximum chemical balancing of reactions, replacing ubiquinone with menaquinone in respiratory chain reactions – *M. tuberculosis* has menaquinone as its only quinone (***Bashiri et al., 2020***) – and introducing a periplasmic compartment to fit the BiGG database identifiers for compatibility with CarveMe. We updated the CarveMe universal bacterial model with the reactions from the revised iEK1011 2.0 model.

We used CarveMe to generate an ensemble of metabolic models for each *Mycobacterium* species by varying the parameters of protein sequence alignment (***Buchfink et al., 2021***), more precisely alignment sensitivity, identity percentage and e-value. The values tested for alignment sensibility were ‘fast’, ‘sensitive’, and ‘more-sensitive’. The values for sequence identity percentage cut-off were 30, 40, 45, 50, 55, 60, and 65, and the values for e-value cut-off 1, 10^−10^, and 10^−50^. The latter two parameters were implemented in the CarveMe code. The reference score used for the reference model was set to 1. We thus generated 63 draft models per species. We checked and corrected, whenever possible, the mass balances of the draft models generated by CarveMe using the corresponding MEMOTE reports (version 0.13.0, ***Lieven et al***. (***2020***)). We ensured that the models satisfy minimal criteria such as network connectivity (removing orphan metabolites), the availability of a chemical formula for each metabolite (completing the model with appropriate formulas from the BiGG database when necessary), and a verified mass balance for every reaction (achieving mass balance for more than 99% of the reactions after correction). We tested the consistency of the candidate models with growth data, as described below. In cases where growth was not predicted by the model but observed in the experiments, we completed the model for this condition using the gapfill function of CarveMe. This was only necessary for *M. smegmatis* in two conditions (L-rhamnose and L-leucine as nitrogen source), however, only L-rhamnose gapfilling was successful.

### Growth predictions using flux balance analysis

Flux balance analysis (FBA) (***Thiele and Palsson, 2010***) was performed on each reconstructed model using the COBRApy toolbox (version 0.29.1) with Gurobi 12.0.3 as linear programming solver (Gurobi Optimization, LLC) (***Schellenberger et al., 2011***). The (lower bound of the) maintenance reaction flux was set to the value of the *M. tuberculosis* model iEK1011 2.0 (3.15 mmol gDW^−1^ h^−1^) (***López-Agudelo et al., 2020***). The maximal uptake rates in the exchange reactions were set to zero, except for oxygen and the components of the growth media (*e*.*g*., water, salts, and trace elements) – Roisin’s minimal medium, modified Sauton’s medium, and Middlebrook 7H9/7H10 media – which were left unconstrained (between 0 and 1000 mmol gDW^−1^ h^−1^). Three types of flux balance analysis were conducted, based on these minimal sets of constraints.

#### Test consistency with growth data during model reconstruction

For testing the consistency of the generated models with the agar plate experiments (**Fig. 2**), the uptake rates of the limiting carbon and nitrogen sources present in the medium were additionally allowed to vary between 0 and 1000 mmol gDW^−1^ h^−1^. In order to predict whether a reconstructed network supports growth on a specific carbon or nitrogen source, FBA was used to maximize flux through the universal bacterial biomass equation (***Xavier et al., 2017***) of CarveMe, where growth corresponds to a flux higher than 0.001 h^−1^. We compared the model predictions over the different conditions with the experimental data by computing the accuracy and Matthew’s correlation coefficient (MCC):

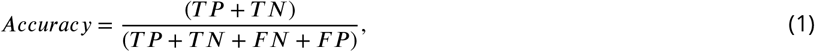

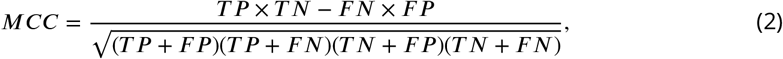

where TP, TN, FP, and FN correspond to the numbers of true positives, true negatives, false positives, and false negatives, respectively.

#### Test consistency of models with measured uptake, secretion, and growth rates

To verify consistency of the reconstructed models with available measurements of uptake, secretion, and growth rates in *M. tuberculosis, M. smegmatis*, and *M. marinum*, we set the exchange fluxes and the growth rate to their measured values plus/minus the experimental error (**Supplementary Table 2**). The units of the measured uptake and secretion rates are the same as in the metabolic models (mmol gDW^−1^ h^−1^), except for *M. marinum* for which measurements were reported in terms of mmol 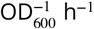(***Dong et al., 2021***). For the latter case, we used a conversion factor from OD_600_ to gDW established in our laboratory (0.39 gDW L^−1^ per unit OD_600_). In order to test the consistency of the models with the constraints imposed by the data, we performed flux minimization by means of parsimonious flux balance analysis (pFBA) (***Lewis et al., 2010***). pFBA successfully returned a solution for all conditions (**Supplementary Table 2**), demonstrating the consistency of the models with the measured uptake, secretion, and growth rates.

#### Predict growth in selected environments

We predicted growth of the five mycobacterial species on 12 carbon sources by means of classical FBA (***Thiele and Palsson, 2010***), setting the uptake rates of the limiting carbon sources to a positive value and maximizing the flux through the biomass reaction. We used knowledge of the protocol of the agar plate experiments and mycobacterial physiology to refine the values for the maximum uptake rate of the carbon source and the cut-off for predicted growth. From the estimated number of bacteria spotted in the beginning of the experiment and the estimated number of bacteria in a visible colony, we determined for each species the minimal growth rate necessary for colonies to appear over the duration of the experiment (0.047 h^−1^ for *M. smegmatis*, 0.027 h^−1^ for *M. aromaticivorans* and *M. abscessus*, 0.013 h^−1^ for *M. marinum* and *M. tuberculosis*). Values of the growth rate above these cut-offs were taken as predictions of growth. Given an approximate value for the biomass yield, in terms of gram dry weight of biomass produced per gram of limiting carbon source, typically around 0.25 for mycobacteria (***Beste et al., 2011; Cook et al., 2009***), an order-of-magnitude for the maximum uptake rate can be obtained by multiplying this value with the growth-rate cut-off. We set the maximum uptake rate within this range and verified that changing its value two- to ten-fold did not much affect the predictions. The use of more refined estimates for the maximum uptake rate and the growth-rate cut-off avoids physiologically unrealistic predictions and thus reduces the number of false positives.

### Functional annotation and statistical tests

We annotated model reactions with top, first- and second-level pathway categories from the KEGG Orthology (KO) database (https://www.genome.jp/kegg/pathway.html, (***Kanehisa et al., 2016***)). Some reactions in the BiGG database were already assigned a KO identifier, for which (after verification) we retrieved the corresponding categories in the KO database. We manually assigned KO identifiers to the remaining reactions by matching model reactions with similar reactions in the KO database (same gene ID, same EC number, or same biochemical reaction). This concerned approximately 1650 reactions, more than 50% of the total. Where appropriate, we created new (second-level) subcategories to classify reactions that had not been previously categorized. By convention, reactions belonging to multiple pathways in the KO database were assigned to a single category.

We defined the core metabolic network as consisting of the reactions shared by all species and the accessory network as consisting of all other reactions occurring in at least one of the species. We tested the over-representation of a given category in the accessory metabolic network as compared to the core metabolic network (**Fig. 3**). For this, we performed one-sided Fisher’s exact tests at the 5% significance level for each functional category. We corrected for multiple testing using the Benjamini-Hochberg procedure (***Benjamini and Hochberg, 1995***).

## Data and software availability

The model reconstruction and analysis pipeline used was implemented in Python 3 and is summarized in **Supplementary Fig. 5**. The code and the user documentation are available from https://gitlab.inria.fr/microcosme/project-myco. The main figures in the paper are automatically generated by the code. The code includes the modified BiGG database, the revised iEK1011 2.0 model for *M. tuberculosis*, and the models generated for *M. aromaticivorans, M. abscessus, M. marinum, M. smegmatis* (with their annotated reactions). The models (in SBML format) and the revised BiGG database are also provided separately (**Supplementary Files 1 and 2**).

## Acknowledgments

The authors acknowledge financial support from the Inria Équipes-associées program (GERM project). They would also like to thank Arnaud Belcour and Michael Baumgärtner for their comments on the manuscript.

